# Equatorial to Polar genomic variability of the microalgae *Bathycoccus prasinos*

**DOI:** 10.1101/2021.07.13.452163

**Authors:** Jade Leconte, Youri Timsit, Tom O. Delmont, Magali Lescot, Gwenael Piganeau, Patrick Wincker, Olivier Jaillon

## Abstract

Phytoplankton plays a fundamental role in the ecology of ocean systems and is the key player in the global carbon cycle. At a time of global warming, understanding the mechanisms of its adaptation to temperature is therefore of paramount importance. Cosmopolitan planktonic species abundant in different marine environments provide both a unique opportunity and an efficient methodological tool to study the genomic bases of their adaptation. This is the case for the eukaryotic picoalga *Bathycoccus prasinos*, whose genomic variability we chose to study in temperate and polar oceanic waters. Using multiple metagenomic datasets, we found that ~5% of *B. prasinos* genomic positions are variable, with an overwhelming majority of biallelic motifs. Cold and temperate waters are clearly associated with changes in variant frequencies, whereas in transitional waters we found more balanced polymorphism at most of these positions. Mesophilic and psychrophilic gene variants are distinguished by only a few amino acid changes located at positions critical for physical and functional protein properties. These results provide new information on the genomic diversity of a cosmopolitan eukaryotic planktonic specie and reveal “minimal mutational strategies” which finely tune the properties of specific proteins at different temperatures.

## Introduction

Protists represent the majority of eukaryotic diversity^1^ and numerous studies address their astounding diversity within marine plankton^2–4^. While eukaryotic diversity among plankton is apparently extremely large, with more than 100,000 species^4^, early studies of genomic diversity tended to indicate some paradoxes. For example, a relatively high rate of interspecific divergence, 70 to 78% amino-acid identity between orthologous proteins, was reported for species of picoalgae within the genera *Ostreococcus*^5^ and *Bathycoccus*^6^. Similarly, while population sizes are expected to be gigantic, the first intraspecific diversity estimations for a variety of eukaryotic phytoplankton species revealed synonymous diversity (*θ_s_*) of around 0.01^7,8^ to 0.02^9^, on par with values measured for multicellular organisms expected to have much lower effective population sizes. In sharp contrast, initial measurements of genetic diversity in the species *Emiliania huxleyi* have shown much lower rates than expected^10^, in seeming contradiction with estimations of mutation rate^11^, highlighting the *Lewontin paradox*^12^.

Until now, a few studies have investigated the spontaneous mutation rate from cultures of mutation accumulation lines, in diatoms^13^, in *Chlamydomonas*^14^ and among the three genera of Mamiellales^15^. The product of the spontaneous mutation rate (*μ*) by the effective population size equals the intraspecific neutral diversity: *θ_s_*= 2*N_e_μ* for haploids, so that both mutation rates and levels of intraspecific diversity are required to infer *Ne*. However, descriptions of intraspecific genetic diversity of eukaryotic plankton populations are scarce as a consequence of the difficulties of cultivation. Other studies presented population genomics analyses from strains within the same species and from multiple sampling sites such as for *Emiliania huxleyi*^10^ or for the Mamiellale *Ostreococcus tauri*^7^, the diatom *Phaedactylum tricornutum*^8^ or the benthic diatom *Seminavis robusta*^9^. Recent studies also addressed the challenge of estimating intraspecific genetic diversity from metagenomes, for example in crustaceans^16^ or marine bacteria^17^

Among phytoplankton, Mamiellales are the most prevalent photosynthetic picoeukaryotes^2,18^. Particularly abundant in coastal waters^19^, they are also widely distributed in open ocean and their geographical distribution has been studied in recent years on the basis of metabarcoding^20,21^ and metagenomics datasets^6,22^. The three main genera of this order, *Bathycoccus*, *Micromonas* and *Ostreococcus*, are distributed over most latitudes and are therefore found in a wide range of environmental conditions^20,22^. However, more precise environmental preferences and biogeographical distribution seem to segregate species within the three genera; *Bathycoccus prasinos*, *Ostreococcus lucimarinus* and *Micromonas pusilla* have been found at significantly lower temperatures than *Bathycoccus* TOSAG39-1 (affiliated to *B. cadidus*^23^), *Ostreococcus* RCC809 and *Micromonas commoda*^22^.

In the Arctic Ocean, which is colder and richer in nutrients than temperate waters^24^, *Micromonas polaris* is largely dominant and seems restricted to this environment^25,26^, however *Bathyccocus prasinos,* which is found in most oceans, is also abundant^27,28^, suggesting an extraordinarily broad distribution of this species. The third genera, *Ostreococcus*, has never been reported in the Arctic despite being present in adjacent seasonally ice-covered waters such as the Baltic sea^29^ or the White sea^30^. In Antarctic waters, only *Micromonas* has been detected, with populations defined as highly similar to the Arctic^31^ ones.

The stringent and abundant detection of the *Bathycoccus* genome in polar, temperate and tropical waters makes this organism a model of choice for analyzing intraspecific genomic diversity, particularly Single Nucleotide Variants (SNVs), in relation to this natural environmental gradient. The evolutionary strategies of cold-adapted organisms are starting to be better understood thanks to the study of the cold-evolved enzymes they are able to produce. This adaptation process can in some instances be related to protein structural changes affecting stability and flexibility^32,33^. Studying *Bathycoccus* structural protein variants *in situ* at different temperatures could further improve our knowledge concerning this mechanism.

Here we leverage metagenomic data of plankton from the *Tara* Oceans collection^34,35^, to compare the genomic diversity of *Bathycoccus prasinos* in polar and temperate environments.

## Materials and Methods

### Genomic resources

*Bathycoccus* RCC1105 was isolated in the bay of Banyuls-sur-mer at the SOLA station at a depth of 3m in January 2006^36^. Sequences were downloaded from the Online Resource for Community Annotation of Eukaryotes^37^. Metagenomics reads from *Tara* Oceans samples^38,39^ corresponding to the 0.8 to 5μm organism size fraction^40^ collected at surface and deep chlorophyll maximum layers of the water column were used to assess the diversity of *Bathycoccus*. For the arctic samples, from TARA_155 to TARA_210, as this size fraction was not available the 0.8 to 2000 μm size fraction was used instead. In stations where both 0.8-5μm and 0.8-2000μm size fraction samples were available we obtained similar *Bathycoccus* relative abundance values (Supplementary Figure 1) probably due to the higher abundance of smaller organisms in plankton.

### Environmental parameters

To assess the potential correlation between genomic variations and local environmental conditions, we used the physicochemical parameter values related to the Tara Oceans expedition sampling sites available in the PANGAEA database^40^. Those contextual data tables can be downloaded at the following link: https://doi.pangaea.de/10.1594/PANGAEA.875582.

### Abundance counts

We mapped metagenomics reads on RCC1105 genome sequences using the Bowtie2 2.1.0 aligner with default parameters^41^. We then filtered out alignments corresponding to low complexity regions with the DUST algorithm^42^ and selected reads with at least 95% identity and more than 30% high complexity bases.

Some gene sequences might be highly similar to orthologous genes, in particular, *Bathycoccus* TOSAG39-1^6^ co-occurring with *Bathycoccus prasinos* in some samples, and thus recruit metagenomic reads from different species. To exclude interspecific mapped reads, we used a statistical approach to discriminate genes with atypical mapping counts. This analysis is based on the assumption that the values of the metagenomics RPKM (number of mapped reads per gene per kb per million of mapped reads) follow a normal distribution. We conducted the Grubbs test for outliers to provide for each sample a list of genes with RPKM distant from this distribution then merged all lists to have a global outliers set^43^. We finally computed relative genomic abundances as the number of reads mapped onto non-outlier genes normalized by the total number of reads sequenced for each sample.

### Filtering steps

Using the previously filtered set of reads, we discarded those with MAPQ scores < 2 in order to remove reads mapping at multiple locations with the same score, which are randomly assigned at either position by Bowtie2 and could cause errors in variant detection. We then calculated genome coverage at each position in the coding regions using BEDTools 2.26.1^44^ and kept samples having an average coverage above 4x. On the initial set of 162 samples, 27 passed this filter. Among them, 4 samples considered to have very high coverage (more than 30x) were selected for a first in-depth variant analysis. The larger set of 27 samples was subsequently used for a global biogeography study.

For each sample, the “callable sites” used in variant analysis were selected from samtools mpileup results^45^ as genomic positions covered by a number of reads comprised between 4 and a maximum corresponding to the average coverage in the sample plus twice the standard deviation.

### Variant calling

We detected variable genomic sites using Anvi’o^46^ on the two sets of *Tara* Oceans samples: a set of four samples above 30x and a set of 27 samples above 4x coverage. For that we created two Anvi’o databases then performed two separate SNV (Single Nucleotide Variant) and SAAV (Single Amino Acid Variant) calls. Quince mode was used in order to retrieve information for each variant locus in all samples. This method takes into account multiple variants in a codon by indicating all amino acids present at a given position rather than independently projecting SNV results. Only positions callable in every sample of interest were kept, in order to make comparisons. For the 4-sample set, a total of 10 585 350 positions (86% of *Bathycoccus* coding regions) were analyzed, while only 1 715 482 positions (14%) were kept across the 27-sample set. We considered an allele to be a variant if it was confirmed by at least 4 reads. Variants were then considered either fixed in a sample (called fixed mutation) if presenting a single allele different from at least one other sample, or polymorphic (called SNV) if presenting two or more alleles within the same sample. Only amino-acid variants for which a corresponding nucleotide variant passed those filters were kept.

### Genomic distance computation

We computed a genomic distance for each pair of samples based on allele content at each SNV position. An allele corresponds here to a nucleotide which can be considered either present or absent, without taking its frequency into account. The distance at each position thus corresponds to the number of common alleles between the samples divided by the sum of the number different alleles in each sample. Identical allelic content would give a score of 1, no allele in common would give a score of 0. The global distance is the average for all positions.

Based on this distance metric, we computed a phylogenetic tree of all samples plus the reference genome RCC1105 using the core R function hclust with default parameters. We obtained bootstrapped values using the pvclust function with 9999 permutations, from the pvclust 2.0-0 R-package. Dendrograms were plotted using the dendextend 1.3.0 package.

Finally, in order to better visualize the information, we used the same values to assign colors to each sample, with the distance between colors reflecting the genomic distance between samples. To achieve this, we carried out Principal Component Analysis (PCA) with package vegan 2.4-1, and translated position values from the three first axes to a Red Green Blue (RGB) color-code for each sample. The resulting color circles were plotted on a map using R-packages ggplot2_2.2.1, scales_0.4.1 and maps_3.1.1.

### Statistical approaches

Multiple statistical analyses were performed for this manuscript based on SNV and SAAV results. First, we computed pairwise water temperature distances in order to run a Mantel test against the genomic distances of all 27 samples, using R-package vegan 2.4-1.

Another experiment focused on variants significantly associated to the large distance between two main groups defined from the hierarchical clusters previously computed. We thus gathered pairwise distances between samples for each position. Then, we selected loci with a mean distance above 0.6 (3.4%) between temperate and cold samples and plotted the density curves of their frequencies for each sample independently using ggplot2_2.2.1.

Finally, we studied the correlation between amino-acid frequencies and sample temperatures in order to assess a potential swap of major and minor alleles between cold and temperate samples. For each SAAV, we took the amino-acid with the highest frequency at the position for each sample, and only kept the positions with at least two different alleles among samples. We then computed a Wilcoxon test for each position using as a first group the temperatures for which the first amino acid was found and as a second group the temperatures for which the other amino acid was found, and applied a Bonferroni correction to the resulting p-values. We kept positions with p-values smaller than 0.05, for a total of 13 variants, and plotted their amino-acid frequencies in our samples using R-packages gridExtra 2.2.1 and ggplot2_2.2.1.

### Protein Homology modeling

The protein sequences of Figure 4 were systematically scanned against the structural databases PDB ^47^ and CATH^48^ to search for structural representatives. Two of them, *Bathycoccus* EPSPS (PDB_id: 5xwb) and eEF3 (PDB_id: 2ix3), displayed sufficient similarity to build reliable models and to locate the sequence variants with respect to their 3D structure. Models of *Bathycoccus prasinos* eEF3, EPSPS and of mesophilic and psychrophilic EPSPS sequences found in the Ocean Gene Atlas web-server (OGA^49^) were constructed using the SWISS MODEL server^50^. The alignments were visualised with JALVIEW ^51^ and the structures were compared with the *Pymol* program^52^. Electrostatic potentials were generated by the vacuum electrostatic function of *Pymol* to obtain a qualitative view of the surface potentials. *Pymol* scripts were used for the systematic localisation of the mutants in the model structures and to quantify and select by distance criterion, mutations of charged residues in the vicinity of pre-existing like-charged residues.

### Search for EPSPS homologs in OGA

The protein sequence of gene Bathy12g01190 potentially encoding a 3-phosphoshikimate 1-carboxyvinyltransferase (uniprot identifier : K8FBR9), involved in chorismate biosynthesis, was used to search for similar sequences in the Marine Atlas of *Tara* Ocean Unigenes (MATOU) with the Ocean Gene Atlas web-server^49^. A total of 1679 eukaryotic metagenomic genes were identified using the default parameters (blastp with a threshold of 1E-10). The selected sequences were separated in two groups based on their strict occurrence within latitude ranges (Mesophilic and Psychrophilic). A total of 227 gene sequences were found only in the Arctic *Tara* oceans stations (from 155 to 210) and named as “arc” (for Arctic). The other gene sequences found in all other latitudes and not in the Arctic stations were named “na” (for not Arctic) and this group contains 678 sequences. The translated metagenomic sequences which aligned with Bathy12g01190 over at least 200 aa were kept. A total of 78 Arctic sequences and 226 non-Arctic sequences were selected and aligned with MAFFT ^53^. The multiple sequence alignments used for phylogenetic analyses contained 631 positions (65 sequences). The best fitting substitution model and rate variation parameters were selected using ProtTest 3^54^ according to the smallest Akaike Information Criterion: WAG+I+G). The phylogenetic reconstructions were performed using PhyML 3.0^55^. Bootstrap values were calculated with 100 bootstrap replicates. The resulting phylogenetic trees were edited using FigTree^56^. The taxonomy of the metagenomic sequences was added to their name in the tree, Supplementary Figure 7.

## Results

### Bathycoccus genomic diversity

We analysed the natural genomic diversity of *Bathycoccus* RCC1105 by mapping metagenomics reads from the *Tara* Oceans expedition^38,39^ on the reference genome^36^ as previously reported^6,22^. Except for chromosomes 14 and 19, known as “outlier chromosomes” (see below), the coverage of recruited reads was relatively homogeneous across genes per sample, with a standard deviation ranging from 30 to 38% of the mean coverage on callable positions (Figure 1).

**Figure 1:**
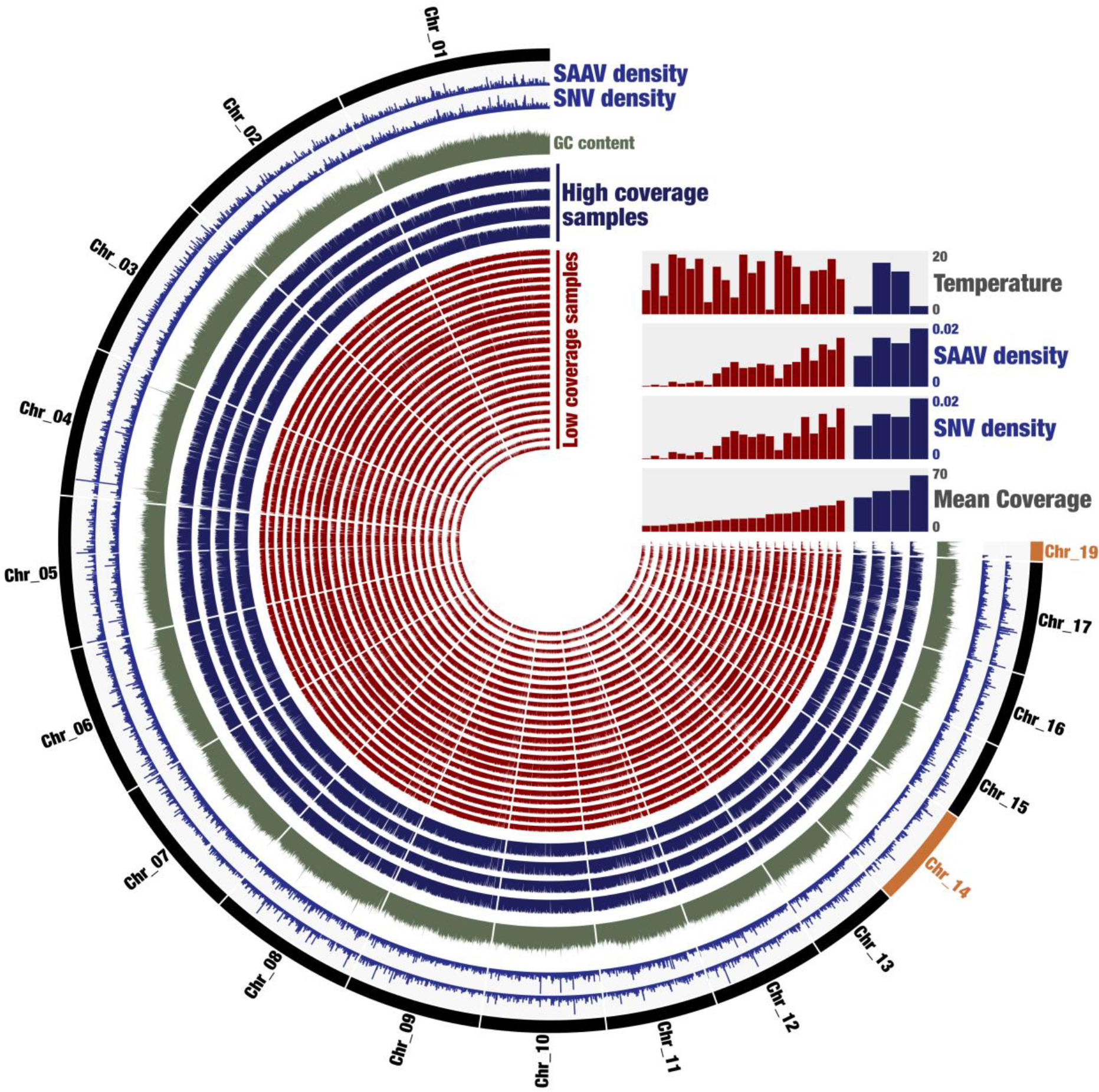
*Bathycoccus* whole genome diversity in 27 Tara Oceans samples. Description of layers starting from the exterior: chromosome number (in black and outliers in orange), SAAV and SNV average density among the four best covered samples (in blue), GC content in percentage (in green), each layer then corresponds to the coverage of one sample, ordered by mean coverage (high-coverage in blue then low-coverage in red). Figure computed with Anvi’o

Chromosome 19, the SOC or Small Outlier Chromosome, has previously been described as hypervariable; accordingly, it presents a major coverage drop along most of its length. Chromosome 14, the BOC or Big Outlier Chromosome, possesses a large region considered as the mating type locus. In our study, unlike the SOC, the coverage is similar to standard chromosomes in the candidate mating type locus of BOC, which would suggest we recruited both mating types. The first third of the chromosome which has a higher GC content presents a slight coverage drop in most samples.

Looking in detail at the Single Nucleotide Variant (SNV) and Single Amino-Acid Variant (SAAV) densities per sample (Figure 1), we observe a positive correlation (spearman test p-value 2.E-07, rho 0.82) between densities and coverage. However, neither densities nor coverage appear to be linked to the water temperature.

For further analysis of the genomic and geographic distribution of SNVs, we applied a minimum threshold of 4X minimum coverage of recruited reads, selecting 27 samples. We also considered a subset of 4 samples where the average coverage of recruited reads is above 30x to detect and possibly quantify the presence of several alleles on more loci. Hence, we identified 11 million genomic positions as callable sites per sample (see methods) on average for the two sets as potential resources for SNVs. We verified the coherence of both sets for comparison of different environments.

Using the reduced set of four samples, a total of 350 478 non-redundant genomic positions present variations in one or more sample, corresponding to 3.31% of our initial set of callable positions (Table 1). The mean coverage goes from 38.41 in one arctic sample to 63.87 in the other one, with two temperate samples falling in between. The total variant density reaches a maximum of 1.96% in the most covered sample, and the majority of those variants correspond to biallelic SNVs.

**Table 1:** Number of variants for the four best-covered samples, total and separation according to the number of alleles found at the loci in each sample.

| Samples | Callable sites | Mean coverage on callable sites | Total number of variants | Number of fixed mutations | Number of biallelic SNVs | Number of triallelic SNVs | Number of quadriallelic SNVs |
| --- | --- | --- | --- | --- | --- | --- | --- |
| 81DCM | 10 972 751 | 45.73 | 146 793<br>(1.38%) | 15 943 | 130 186 | 662 | 2 |
| 135DCM | 10 983 591 | 44.80 | 155 249<br>(1.46%) | 14 310 | 140 083 | 854 | 2 |
| 196SUR | 11 181 235 | 62.13 | 208 320<br>(1.96%) | 7 557 | 199 024 | 1732 | 7 |
| 209SUR | 11 023 554 | 37.77 | 108 967<br>(1.03%) | 14 303 | 94 463 | 201 | 0 |

The coverage and number of callable sites are consistent between chromosomes for all samples, except for chromosome 19 (79 to 91% against 23% for callable sites) which has a very low horizontal and vertical coverage (Supplementary Table 1). The variant density appears homogeneous among chromosomes, except for chromosome 14, where the density drops to half of that found for the other chromosomes. Variant density also appears homogeneous across the four different samples (Supplementary Figure 2).

At the codon level, we detected SNVs leading to amino-acid changes (SAAVs, as previously reported^17^) to approximate the ratio of non-synonymous versus synonymous variants by dividing the number of SAAVs by the number of SNVs. About 6% of codons containing SNVs have more than one nucleotide variant. Here, the SAAV/SNV ratios per sample range from 0.33 to 0.40 without apparent geographical or environmental patterns for the four samples. By associating each SAAV with an amino acid substitution type (AAST), defined as the two most frequent amino acids in a given SAAV, we obtained a distribution of observed AAST frequencies^57^ (Supplementary Figure 3) similar to the one found with the same rationale for a bacterial population detected at higher abundance^17^. This confirms the validity of our dataset for further analyses of SNVs and SAAVs.

On the larger set of 27 metagenomic samples, the density of variants was positively linked among the sets with an almost perfect linear correlation (Supplementary Figure 4) and SAAV/SNV ratios are also similar, validating use of this larger set of samples on a reduced portion of the genome. In these 27 samples we obtained a total of 80 284 SNVs and fixed mutations which correspond to 4.68% of callable positions (Supplementary Table 2). This higher number of variant alleles compared to the previous value on four samples is expected given the greater sample size. As previously observed, most samples present more polymorphic positions than fixed allele positions among our set of callable sites. A notable exception, samples from southern waters provided a very low SNV rate for good read coverage. At the SAAV level, we cannot see any particular correlation between the SAAV/SNV ratio and either environmental conditions or geographical patterns at oceanic basin scale. AAST distributions are almost identical using the 4 and 27 sample sets, with only minor inversions in AAST prevalence.

### Population structure analysis

To compare populations at SNV level, we computed pairwise genomic distances between all samples considering fixed and polymorphic positions within the set of nucleotide variations found in the 27 samples (Methods). A dendrogram representing these distances among samples clearly separates Arctic and temperate samples (Figure 2). The southern samples (numbers 82 and 89) are the furthest away from all the other groups, perhaps related to their particular environment, but even more surprisingly, the Mediterranean Sea sample 9DCM is also positioned away from the others. Those three samples present much lower polymorphism.

**Figure 2:**
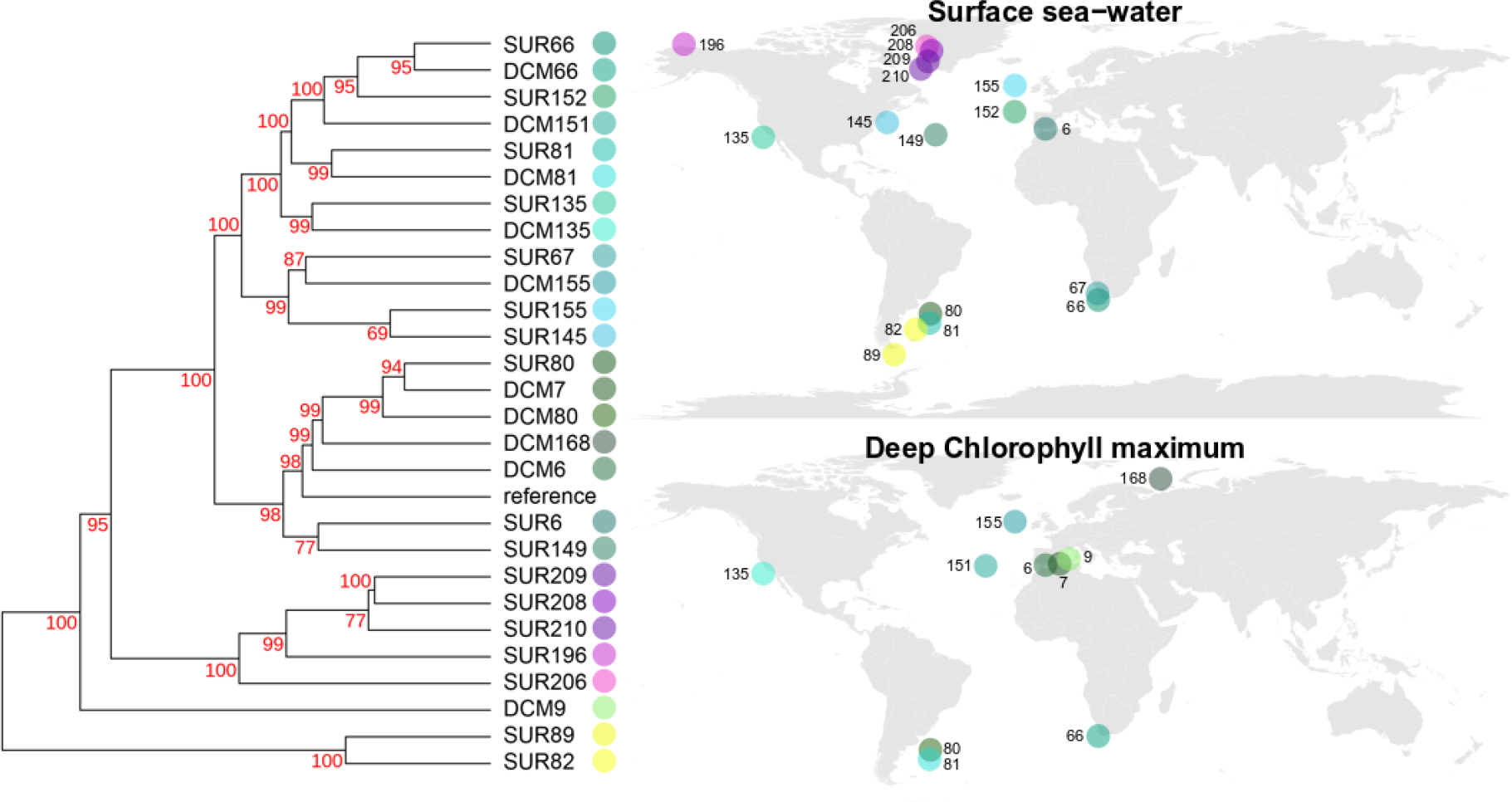
*Bathycoccus prasinos* genomic distance among 27 sampled populations based on nucleotide variant patterns, displayed as a phylogenetic tree (left panel) and a biogeography map (right panel for surface (top) or Deep Chlorophyll Maximum (bottom)). The similarities of sample colors (right) reflect the genomic similarities of populations (left) (Methods).

To facilitate visual interpretation of a geographic representation of these genomic distances, each population was assigned a color such that the difference in color between populations reflects their genomic distance (Methods, Figure 2). Multiple biogeographical patterns emerged, for example a clear separation of Arctic samples in purple and austral ones in yellow. Most temperate samples are represented in a gradient of green, but sample 145SUR, near the end of the Labrador current thus under the influence of Arctic waters, and 155SUR near the end of the Gulf Stream just before Arctic waters, are both represented in blue. Sample 135DCM, located off the coast of California near a site of cold and rich upwelling and samples 81SUR and 81DCM situated close to austral waters are represented in blue-green colors, intermediate between temperate and warm-cold transition points. The Mediterranean sample 9DCM mentioned above appears in a yellow-green color, which is coherent with its genomic patterns less distant from the austral samples.

A Mantel test between matrices of genomic and temperature differences confirms a positive correlation (0.4818 statistic at p-value=0.001, Supplementary Figure 5).

### Genomic differences between temperate and cold-water populations

We identified a set of 2742 SNVs that significantly distinguish temperate and cold populations (Methods) and verified very different allelic frequency distributions (Figure 3). Using the complete set of SNVs, we retrieved similarly shaped allelic frequency distributions for temperate populations but not for cold ones (Supplementary Figure 6). In temperate samples, a majority of alleles are fixed, observed frequencies are close to 0% or to 100%. We cannot rule out the possibility that other variants are present in populations at rates below our detection capacity. In contrast, in all but one of our arctic samples (sample 206), the genomic positions of this set mostly present local polymorphisms between two principal alleles, present in different ratios depending on the sample (80/20, 70/30 or 50/50 ratios). The high genomic distances between arctic and temperate populations would thus be mainly related to this group of loci that appear biallelic in the Arctic but present a single allele elsewhere.

**Figure 3:**
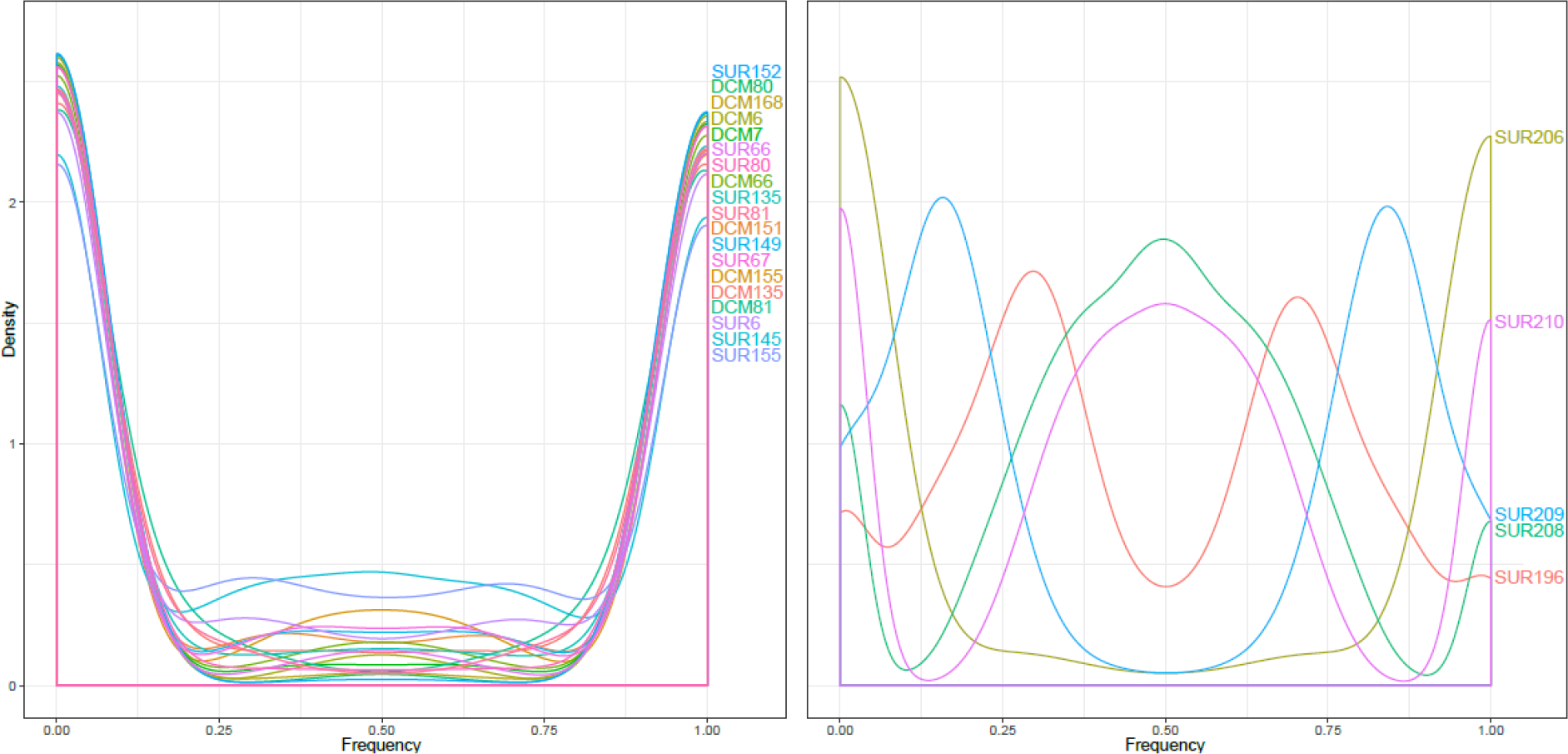
Allele frequency distributions for 2742 selected SNVs for different samples belonging to the temperate (left panel) or the arctic (right panel) cluster.

Finally, we examined amino-acid changes between cold and temperate populations by selecting SAAVs for which each allele was significantly statistically associated with a different water temperature (Methods), resulting in a set of 13 genomic positions. We analyzed their allele frequencies with respect to the genes in which they were located (Figure 4) and found a clear pattern of swap of major amino-acids between temperate and cold samples, for which arctic and austral populations have similar phenotypes for those positions. In most samples, the amino-acid frequencies of these positions correspond to fixation or near-fixation. Interestingly, populations presenting a protein polymorphism were often sampled at intermediate temperatures and also at transition points between different oceanographic systems (samples 145 in North-Atlantic or 81 in South-Atlantic).

**Figure 4:**
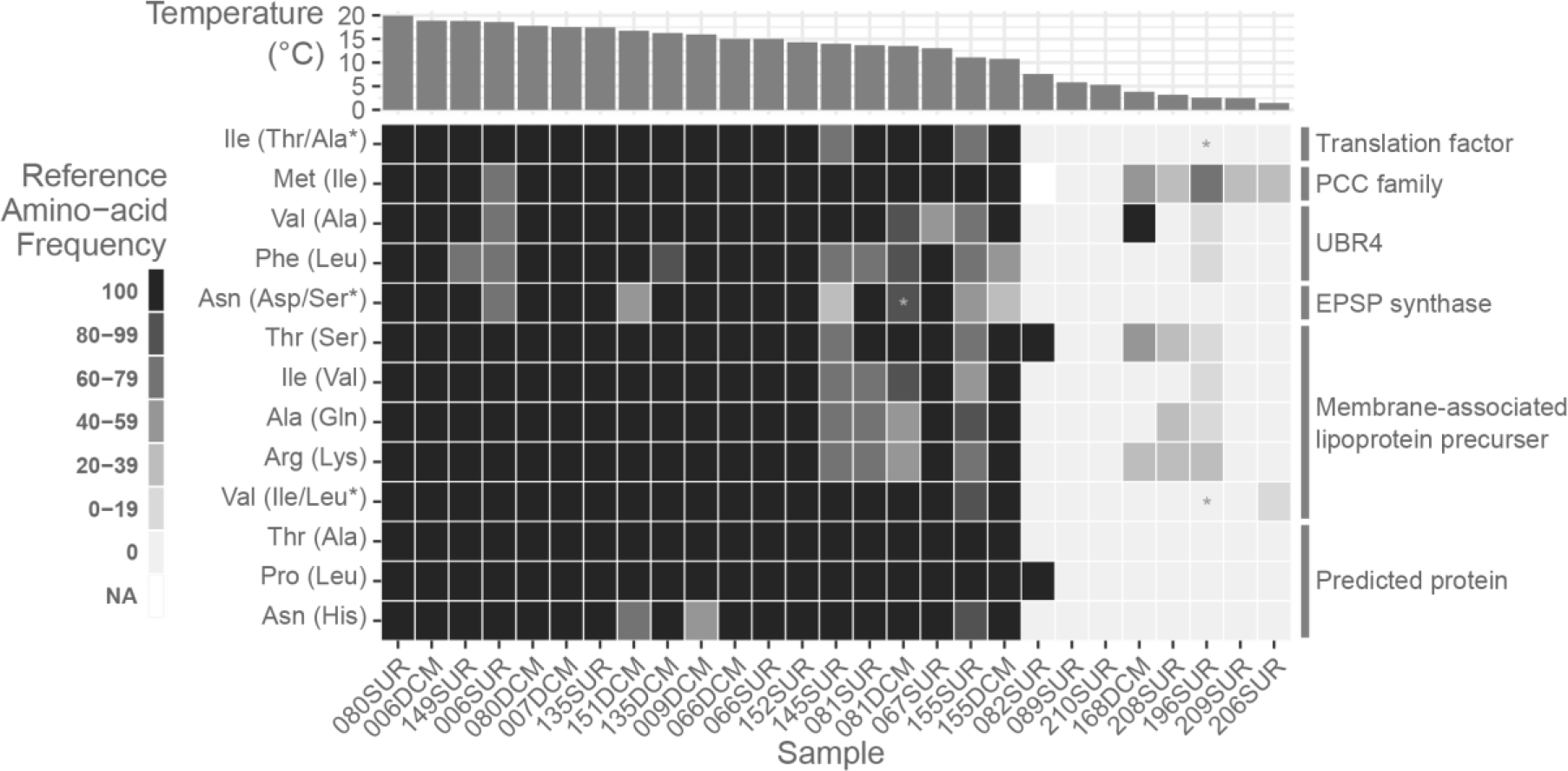
Heatmap presenting the frequency of amino-acids among 27 samples in 13 genomic positions that segregate populations according to temperature. Alternate amino-acids found are indicated in parentheses. Samples are sorted by their temperature indicated at the top, and the six proteins in which variants are found are indicated on the right. From top to bottom, the proteins correspond to Bathy02g02490, Bathy03g01170, Bathy09g03590, Bathy12g01190, Bathy13g00360, Bathy13g00290.

In an attempt to elucidate the molecular mechanisms of cold adaptation, we sought to locate the single mutations in the structure of proteins listed in Figure 4. One of them contains three amino-acid mutations but has no functional annotation. Others appear related to stability of the protein structure or to low or high temperature, such as the membrane-associated lipoprotein precursor containing six mutations, or the PCC (Polycystin Cation Channel) protein. Indeed structure and concentration of lipids has been shown to be important for cold adaptation in many organisms^58,59^ and transport proteins might need specific adaptations in order to function at low temperatures^60,61^.

Two sequences have close structural homologues making it possible to build reliable models and locate the mutations. The first one displays 47.26 % identity with the yeast translation factor eEF3 (PDB_id: 2xi3)^62^ that is composed of 5 subdomains (HEAT, 4HB, ABC1, ABC2 and chromo). The I67T mutation is localized in the HEAT domain^63^ consisting of a repeat of 8 pairs of α-helices forming the N-terminal domain (1-321) of the yeast eEF3 structure. However, the alignment of yeast and *Bathycoccus* eEF3 sequences reveals that the yeast sequence possesses an insertion (H43-S77; yeast eEF3 numbering) that corresponds to the second pair of α-helices of the HEAT domain. Thus, the *Bathycoccus* eEF3 HEAT domain has only 7 helix pairs and the I67T mutation is located at the C-ter end of its third α-helix, which contacts the 4HB domain (Figure 5a). The replacement of an Ile with a less hydrophobic Thr likely destabilises the packing of the amphiphilic α-helices on the inner side of the HEAT domain and probably contributes to making it less stable and more flexible.

**Figure 5:**
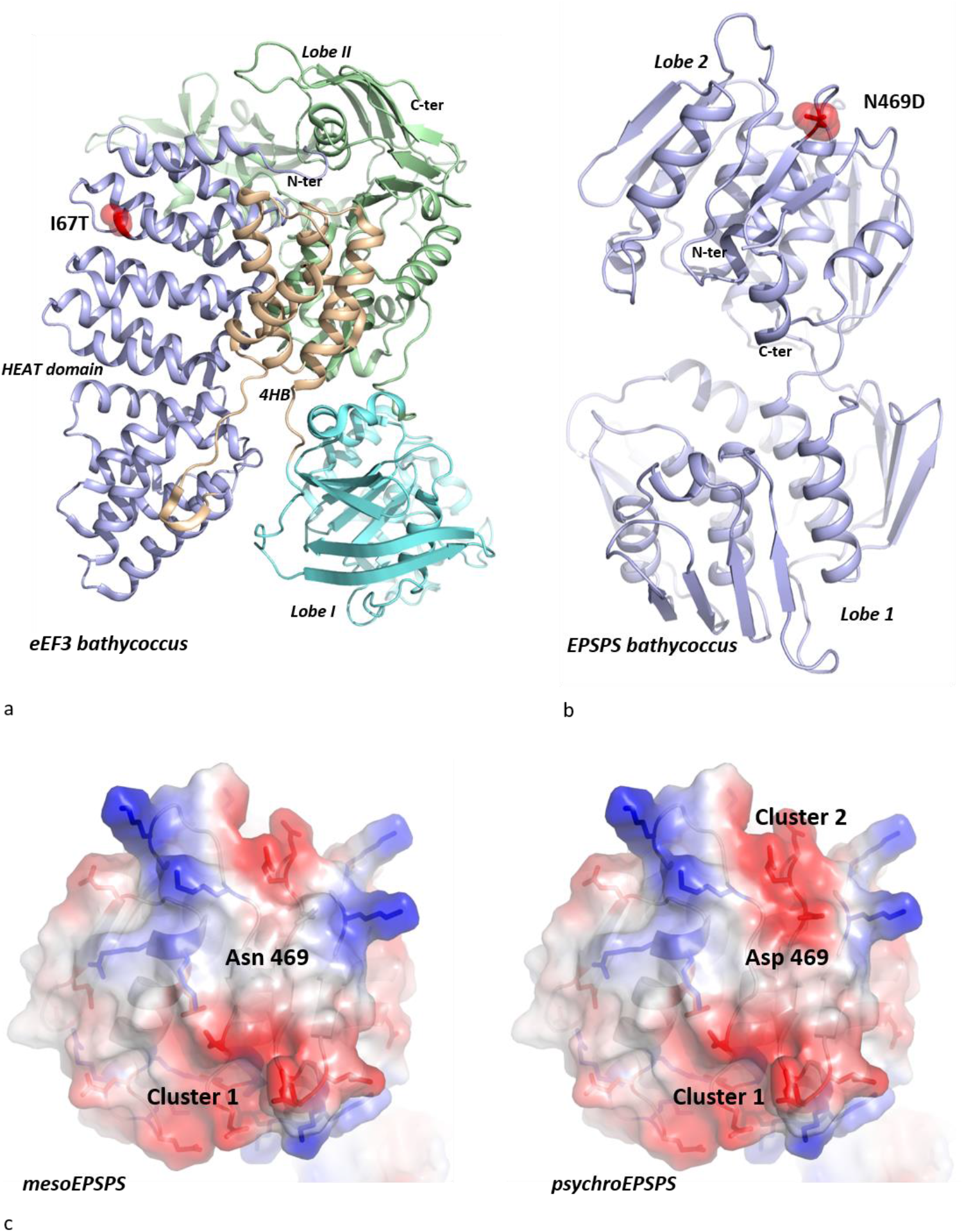
Positions of the single mutations in the structure of eEF3 and EPSPS. a : *Bathycoccus prasinos* eEF3 model. b: *Bathycoccus prasinos* EPSPS model. The locations of the mutations are represented by a red sphere. c: Comparison of the electrostatic surface potentials of the *Bathycoccus prasinos* mesophilic (left) and psychrophilic (right) EPSPS models. Cluster 1 is formed by the grouping of E60, E61, D474, D449 and D452, on the edge of domain 2 of both mesophilic and psychrophilic *Bathycoccus prasinos* EPSPS. Cluster 2 is formed by the grouping of D457, E468 and the mutated D469 in the psychrophilic EPSPS.

The second sequence corresponds to 5-enolpyruvylshikimate-3-phosphate synthase (EPSPS) (PDB_id: 5xwb), the 6th enzyme of the shikimate pathway^64^ that leads to the biosynthesis of aromatic amino acids. This enzyme consists of two lobes, each composed of an arrangement of 3 similar subdomains consisting of a βαβαββ fold^65^ (Figure 5b). The comparison of the *Bathycoccus prasinos* EPSPS model to its structural homologs in the PDB indicates that this enzyme belongs to the glyphosate-insensitive EPSPS class II^66^. It does not possess the highly conserved ‘90-LFLGNAGTAMRPLAA-104’ motif characteristic of class I, and it has the RPMxR motif that has been shown to be responsible for the glyphosate insensitivity of class II enzymes^67,68^. This is an interesting observation as it was thought that only bacterial EPSPS had class II representatives. The N469D mutation is located on the surface of domain 2, on a loop joining the α13-helix and the β23-sheet. It corresponds to proline 401 of its closest homolog, EPSPS from *Colwellia Psychrerythreae* (54.82 % identity)^69^. Interestingly, in the psychrophilic protein, the N469D mutation appears right next to two other acidic amino acids (E468, D467) and generates a cluster of negative charges on its surface (cluster 2) (Figure 5c). This coulombic repulsion may therefore contribute to destabilize the protein surface of the cold variant. However, another particular feature of this variant may also help to explain its critical role in cold adaptation. The newly formed cluster 2 arises in the vicinity (17Å) of another negatively charged cluster (cluster 1) shared by both the mesophilic and psychrophilic *Bathycoccus* EPSPS forms (Figure 5c). It is therefore likely that this proximity contributes further to destabilize the psychrophilic protein. This illustrates how a single but strategically poised mutation can produce a global effect on the overall electrostatic properties and thus the stability of the protein. This provides a “minimal” evolutionary strategy that exploits contingently the protein surface features to tune its stability with single mutational events. To test the generality of this evolutionary mechanism, we extended our analysis to eukaryotic EPSPS homologs found in the *Tara Oceans* metagenomic data.

Using the OGA server^49^, we selected two groups of sequences distributed specifically according to their latitude, one strictly confined to the poles and the other strictly present in mesophilic areas. The sequences of these two groups were then aligned together and a tree was constructed to select the closest pairs of psychrophilic and mesophilic planktonic eukaryotic EPSPS (Supplementary Figure 7). The comparison of each sequence pair identified a set of mutations most likely associated with psychrophilic life and allowed their localisation in the corresponding structures. The vast majority of mutations occurs on loops at the surface of the two domains and are distributed on equivalent structural motifs through the pseudo 3-fold symmetry of each lobe (Supplementary Figure 8). Interestingly, among the mutations that involve charged residues (34% of the total mutations, Supplementary Figure 9 a-c), 86% of the acidic and 62% of the basic mutated amino acids, respectively, appear in close proximity of like-charged amino acids (Supplementary Figure 9d). Positively charged clusters are also observed in cold adaptation in this enzyme family (Supplementary Figure 9e). This analysis therefore supports the idea that creating clusters of like-charged residues on the EPSPS protein surface constitutes an evolutionary strategy for cold adaptation.

## Discussion

Using metagenomics samples from the *Tara* Oceans expedition and the genome sequence of the picoeukaryote alga *Bathycoccus prasinos* RCC1105, we assessed the genomic diversity of a cosmopolitan species model in temperate and polar marine biomes. With 27 metagenomic samples where this genome presents a significant coverage of recruited reads, we estimated that single nucleotide variations are mostly present in biallelic forms with a maximum density reaching 2% of coding regions. Polar and temperate populations appear to be clearly segregated by single nucleotide variations in 0.16% of the genomic positions. Thus, we presented a clustering of *Bathycoccus* populations segregating into three groups based on those nucleotide variations, corresponding to austral, arctic and temperate waters, and mainly characterized by the presence or absence of polymorphism on the segregating variants.

We were able to detect six proteins presenting a highly biome-dependent amino acid composition. Cold-adapted proteins are generally more flexible either around the active site or on their surface to optimize their functionality at low temperatures^32,33,70–72^. However, the evolutionary mechanisms by which a single nucleotide variant alters the global properties of a protein remain poorly understood. A recurrent obstacle is that very often the compared mesophilic and psychrophilic proteins belong to different species and have greatly diverged, making it difficult to discern nucleotide variations that are strictly related to temperature adaptation. Our paper presents an opportunity to circumvent this obstacle since it compares proteins of the same ubiquitous species, with mesophilic and psychrophilic variants, thus highlighting nucleotide and amino-acid variations specifically linked to habitat changes. *Bathycoccus* EPSPS and eEF3 show two distinct modes of cold adaptation. While in eEF3 it is the reduction of the hydrophobic character that alters the stability of an enzyme domain, it is through the modification of the electrostatic surface properties that EPSPS adapts to cold. However, in both cases, the evolutionary strategy is remarkable in the sense that a single amino-acid variation was sufficient to optimise the properties of the enzyme at low temperatures. We show that these mutations occupy critical positions in protein structures whose modification induces global changes of their physical and functional properties.

According to our results the N469D amino-acid variant found in the psychrophilic *Bathycoccus* EPSPS produces a cluster of negative charges on the protein surface, in close proximity to a pre-existing cluster of the same charge. The resulting coulombic repulsions at the cold-adapted *Bathycoccus* EPSPS surface likely enhance the flexibility that is generally required for keeping functionality at low temperatures. Interestingly, the I67T amino acid variant found in the psychrophilic *Bathycoccus* variant is located on the third helix of the HEAT domain of eEF3. This reflects a highly targeted evolutionary pressure on its HEAT domain and suggests that the level of flexibility of one of its HEAT repeats is essential for its function. The key to understanding the mechanistic importance of this mutation lies in the physical properties of HEAT repeats^63,73^. HEAT domains possess remarkable elastic properties that allow them to reversibly undergo multiple mechanical stresses^74–76^. The elastic properties of HEAT repeats rely on an unusual hydrophobic core that differs significantly from that of less flexible globular proteins^77^. Moreover, it has been found that the HEAT domain function depends on a non-uniform distribution and of the stability of each repeat within the HEAT domain^76^. Knowing that the replacement of an isoleucine by a threonine contributes to the cold adaptation adenylate cyclase by altering the packing of its hydrophobic core^78^, the present study documents how the subtle adjustment of the hydrophobic properties of the 2^nd^ HEAT repeat adapts its elasticity at different temperatures. To our knowledge, this constitutes the first data suggesting the HEAT repeats may be involved in adaptation to cold temperatures (Supplementary Information).

The North Atlantic Current forms the southern and eastern boundary current of the subpolar gyre circulation, crossing the North Atlantic before flowing into the Iceland Basin^79^. Plankton is passively transported along this path and encounters the polar front; *Bathycoccus prasinos* RCC1105 seems to cross it with success as indicated by the very high genomic similarity between the abundant polar and temperate populations. Therefore, in light of our results indicating a population structure depending on water temperature, multiple hypotheses can be raised concerning the evolutionary strategies that have shaped the genomic properties of *Bathycoccus prasinos*. Among these, the existence of alleles that would be restricted to each biome appears highly unlikely. Indeed, the polymorphic genomic loci of *Bathycoccus prasinos* populations consist mainly of two alleles whose proportions vary along the path of the currents connecting arctic and temperate waters. We favor the hypothesis that a relatively short life cycle combined with environmental selection occurring along the path would permit rapid recombination of dominant alleles and rapid swap of their relative proportions in populations transported by currents.

In line with this proposition, populations sampled in marine areas located between cold and temperate oceans present polymorphisms at those segregating amino-acid positions.

Further studies, from cultures or from natural populations, are required to better characterize the functional impact of these amino-acid variations. For example, the probable adaptation patterns would benefit from a gene expression study to test patterns of acclimation, as recently exemplified in fish^80^ complementing patterns observed for bacterial communities in the same samples^35^. Such data would feed into efforts to better understand and predict the impact of global warming, which could have a major impact on the polar biome community^81^. It has recently been suggested that advection by North Atlantic currents to the Arctic Ocean, combined with warming, will shift the distribution of phytoplankton poleward, leading to a restructuring of biogeography and complete communities^82^. Such rapid and significant changes challenge adaptation and acclimatization strategies that have evolved over millions of years, especially for cosmopolitan organisms with a temperature-related population structure such as *Bathycoccus prasinos*.

This study exploits the opportunity to have sampled some of the natural genomic variability of *Bathycoccus prasinos* populations from different temperate and polar locations. Although this geographic coverage is relatively broad, as is the sequencing effort, it is very likely that we have captured only a small portion of the genomic diversity of this species. However, we do see possible markers of adaptation to the Arctic zone where environmental selection pressure would be exerted on the molecular dynamics of proteins due to low temperatures.

With more samples, future studies will have to consider variations of both genomes and protein molecular dynamics together the with spatio-temporal context of connectivity among plankton communities to uncover evidence of adaptations to other environmental pressures.

## Acknowledgments

We would like to thank all Genophy group and LAGE members for stimulating discussions on this project. Tara Oceans would not exist without the Tara Ocean Foundation and the continuous support of 23 institutes (https://oceans.taraexpeditions.org/). This article is contribution number XX of Tara Oceans.

## Author contributions

P.W. and O.J. designed the study.

J.L., Y.T. and O.J. wrote the paper with contributions of G.P.

J.L. performed genomic diversity and population structure analysis with contributions of T.D., G.P. and O.J.

Y.T. and M.L. performed structural analysis.

All authors have read and agreed to the published version of the manuscript.

