## Supplementary Information for "Equatorial to Polar genomic variability of the microalgae *Bathycoccus prasinos*"

***EPSPS***

We show that the N469D mutation found in the psychrophilic *Bathycoccus* EPSPS produces a cluster of negative charges on the protein surface, in close proximity to a pre-existing cluster of the same charge. By creating contiguity between two negatively charged clusters, this single amino acid change may have a 'critical effect' that can affect the overall properties of the protein. The resulting coulombic repulsions at the cold-adapted *Bathycoccus* EPSPS surface likely enhance its flexibility that is generally required for keeping functionality at low temperatures. Furthermore, our analysis of other eukaryotic planktonic EPSPSs generalizes this evolutionary strategy for the cold adaptation of this enzyme. Of the six sequences analysed, 87% and 62% of mutations that introduce negative or positive residues, respectively, occur in close proximity of residues of the same charge. For example, a psychrophilic *dinophyceae* EPSPS exploits a similar surface destabilization strategy but with positive charges. These observations are consistent with previous works that show the importance of the protein electrostatic in cold adaptation[1–6]. The distribution of charged residues in protein sequences and their electrostatic interactions contribute significantly to protein stability [2,4,7] . Experimental and theoretical studies have shown that mutation of a single amino acid on the surface can alter the overall protein stability [8–10]. For example, elimination of like-charge repulsion and creation of opposite-charge attractions on the protein surface enhance the protein stability. In contrast, a crystal structure that trapped both a folded and unfolded intermediate of a ribosomal protein provided the experimental evidence that clusters of like-charged residues were directly responsible for the unfolding of certain protein regions and that this local instability could have global repercussions on the overall protein structure [11].

***eEF3***

Originally thought to be restricted to the fungi group [12,13], eEF3 has been progressively found in other eukaryotes [14–16]. eEF3 is involved in the translation elongation cycle [13,17] and in ribosome recycling [18]. However, many studies show that in yeast, eEF3 is also involved in functions other than translation such as the post-transcriptional regulation of many mRNAs, protein targeting in the plasma membrane [19] and one of its paralog is essential for growth under oxidative stress[20] . With 870,000 copies per cell[21], which largely exceeds the copy number of eEF1A and of ribosomes[22] , the eEF3 abundance is thought to reflects its pleiotropic role in yeast [19]. Furthermore, eEF3 associates with a Cch1 calcium channel and directs it to the plasma membrane in *Cryptococcus neoformans* [23]. The targeting roles of eEF3 in either binding to the 40S subunit of the ribosome [17] or in regulating the localisation of Cch1 in the plasma membrane [23] depends on its HEAT domain. Interestingly, the HEAT domains of many proteins such as Tor2 [24,25] or Huntingtin [26] also play a key role in cell signalling or protein targeting by interacting with numerous partners. Interestingly, the I67T mutation found in the psychrophilic *Bathycoccus* variant is precisely located on the third helix of the HEAT domain of eEF3. This reflects a highly targeted evolutionary pressure on its HEAT domain and suggests that the fine-tuning of the flexibility of one of its HEAT repeat is essential for its function. This also underlines that the protein property altered by this mutation cannot be circumvented by alternative strategies of adaptation to temperature such as post-transcriptional modifications or the addition of solutes to the cytoplasm[27]. The keys to understanding the mechanistic importance of this mutation lie in the physical properties of HEAT repeats [28,29]. HEAT domains possess remarkable elastic properties that allow them to undergo reversibly multiple mechanical stresses [30–32]. Interestingly, the elastic properties of HEAT repeats rely on an unusual hydrophobic core that differs significantly from those of less flexible globular proteins [33]. Moreover, it has been found that the HEAT domain function depends on a non-uniform distribution and a fine-tuning of the stability of each repeat within the HEAT domain [32]. Knowing that the replacement of an isoleucine by a threonine contributes to the cold adaptation adenelytate cyclase by altering the packing of its hydrophobic core [34], our study thus documents how the subtle adjustment of the hydrophobic properties of the 2$^{nd}$ HEAT repeat adapts its elasticity at different temperatures. To our knowledge, this constitutes the first data that documents the cold adaptation of HEAT repeats. They are not only interesting from an evolutionary point of view but also provide information on the physical and functional properties of a domain of considerable functional importance in cells in which particular mutations may cause cancer[35–37].

| Chr. | Total size | Coding size | Callable sites % | Mean cov. 81DCM | Mean cov. 135DCM | Mean cov. 196SUR | Mean cov. 209SUR | Variant density 81DCM | Variant density 135DCM | Variant density 196SUR | Variant density 209SUR |
|---|---|---|---|---|---|---|---|---|---|---|---|
| 01 | 1 352 724 | 1 133 982 | 91% | 45.7 | 45.63 | 65.14 | 38.57 | 1.31% | 1.41% | 1.95% | 0.97% |
| 02 | 1 122 692 | 922 092 | 90% | 46.16 | 45.54 | 63.86 | 38.31 | 1.38% | 1.48% | 2.02% | 1.03% |
| 03 | 1 091 008 | 909 405 | 89% | 46.65 | 45.65 | 64.02 | 38.29 | 1.36% | 1.43% | 2.02% | 1.02% |
| 04 | 1 037 991 | 863 091 | 87% | 47.82 | 46.54 | 65.36 | 39.65 | 1.35% | 1.38% | 1.91% | 1.04% |
| 05 | 1 019 276 | 850 563 | 89% | 46.95 | 45.81 | 65.17 | 38.78 | 1.39% | 1.48% | 1.91% | 1.03% |
| 06 | 989 707 | 821 307 | 89% | 45.76 | 45.03 | 63.75 | 38.77 | 1.37% | 1.49% | 2.05% | 1.07% |
| 07 | 955 652 | 791 748 | 87% | 47.16 | 46.82 | 65.2 | 39.91 | 1.31% | 1.42% | 1.75% | 0.92% |
| 08 | 937 610 | 767 367 | 89% | 47.08 | 46.31 | 64.27 | 38.95 | 1.30% | 1.39% | 1.91% | 1.02% |
| 09 | 895 536 | 740 394 | 89% | 46.5 | 45.39 | 64.35 | 38.87 | 1.30% | 1.38% | 1.95% | 1.00% |
| 10 | 794 368 | 667 857 | 88% | 46 | 45.38 | 62.7 | 37.78 | 1.40% | 1.45% | 2.05% | 1.09% |
| 11 | 741 603 | 610 035 | 86% | 45.59 | 45.42 | 63.02 | 38.92 | 1.43% | 1.51% | 2.08% | 0.99% |
| 12 | 712 459 | 597 036 | 90% | 46.33 | 45.89 | 63.17 | 38.55 | 1.42% | 1.50% | 2.11% | 1.08% |
| 13 | 708 035 | 607 836 | 79% | 46.79 | 44.95 | 64.57 | 37.37 | 1.52% | 1.51% | 1.97% | 1.19% |
| 14 | 663 424 | 478 431 | 90% | 47.43 | 46.5 | 60.31 | 34.77 | 0.98% | 1.04% | 1.00% | 0.53% |
| 15 | 519 835 | 431 610 | 87% | 46.53 | 45.38 | 64.31 | 38.46 | 1.40% | 1.49% | 1.94% | 1.05% |
| 16 | 494 108 | 407 472 | 84% | 47.33 | 46.14 | 62.55 | 38.19 | 1.43% | 1.50% | 1.98% | 0.99% |
| 17 | 465 570 | 386 298 | 78% | 46.02 | 44.99 | 61.94 | 37.25 | 1.57% | 1.67% | 2.05% | 1.15% |
| 18 | 310 170 | 251 943 | 84% | 44.99 | 43.89 | 60.16 | 36.87 | 1.78% | 1.86% | 2.30% | 1.24% |
| 19 | 146 238 | 84 381 | 23% | 26.46 | 27.17 | 35.01 | 19.29 | 1.41% | 1.50% | 1.16% | 0.62% |
| Total | 14 958 006 | 12 322 848 | 87% | 45.43 | 44.65 | 62.05 | 37.24 | 1.37% | 1.44% | 1.94% | 1.01% |

**Supplementary Table 1:** Main statistics of genomic diversity among populations of *Bathycoccus prasinos* from the four best-covered samples. For each chromosome or the whole genome, the columns indicate total chromosome size, cumulative length of coding regions, percentage of the chromosome considered as callable, coverage and variant density for each of the samples.

| Samples | Callable sites | Mean coverage on callables | Total number of variants | Number of fixed mutations | Number of biallelic SNVs | Number of triallelic SNVs | Number of quadriallelic SNVs |
|---|---|---|---|---|---|---|---|
| 6DCM | 8 804 921 | 8.22 | 8 662 *(0.50%)* | 4 786 | 3 876 | 0 | 0 |
| 6SUR | 10 698 809 | 20.01 | 16 627 *(0.97%)* | 2 491 | 14 109 | 27 | 0 |
| 7DCM | 10 053 998 | 10.94 | 7 624 *(0.44%)* | 3 873 | 3 749 | 2 | 0 |
| 9DCM | 6 355 421 | 6.53 | 13 171 *(0.77%)* | 11 293 | 1 878 | 0 | 0 |
| 66DCM | 10 646 963 | 21.67 | 15 940 *(0.93%)* | 3 662 | 12 261 | 17 | 0 |
| 66SUR | 10 030 196 | 12.45 | 11 829 *(0.69%)* | 4 613 | 7 213 | 3 | 0 |
| 67SUR | 10 172 546 | 14.89 | 16 710 *(0.97%)* | 4 673 | 12 027 | 10 | 0 |
| 80DCM | 9 324 737 | 8.84 | 7 098 *(0.41%)* | 4 084 | 3 012 | 2 | 0 |
| 80SUR | 10 778 634 | 19.82 | 8 005 *(0.47%)* | 3 087 | 4 915 | 3 | 0 |
| 81DCM | 10 972 751 | 45.73 | 26 111 *(1.52%)* | 3 184 | 22 843 | 84 | 0 |
| 81SUR | 10 688 629 | 26.93 | 18 300 *(1.07%)* | 4 431 | 13 846 | 23 | 0 |
| 82SUR | 6 644 362 | 6.48 | 14 599 *(0.85%)* | 14 220 | 379 | 0 | 0 |
| 89SUR | 7 104 067 | 6.98 | 14 322 *(0.83%)* | 13 818 | 504 | 0 | 0 |
| 135DCM | 10 983 591 | 44.8 | 27 574 *(1.61%)* | 3 119 | 24 344 | 111 | 0 |
| 135SUR | 10 597 732 | 28.99 | 20 978 *(1.22%)* | 3 674 | 17 245 | 59 | 0 |
| 145SUR | 10 894 102 | 28.8 | 26 989 *(1.57%)* | 2 461 | 24 462 | 66 | 0 |
| 149SUR | 10 272 274 | 14.3 | 16 645 *(0.97%)* | 3 340 | 13 294 | 11 | 0 |
| 151DCM | 10 197 027 | 15.04 | 17 967 *(1.05%)* | 4 345 | 13 596 | 26 | 0 |
| 152SUR | 9 172 532 | 9.35 | 9 276 *(0.54%)* | 7 231 | 2 044 | 1 | 0 |
| 155DCM | 10 072 636 | 12.96 | 16 263 *(0.95%)* | 4 255 | 11 999 | 9 | 0 |
| 155SUR | 10 983 560 | 34.28 | 29 959 *(1.75%)* | 2 447 | 27 399 | 113 | 0 |
| 168DCM | 10 084 235 | 11.53 | 6 399 *(0.37%)* | 5 364 | 1 035 | 0 | 0 |
| 196SUR | 11 181 235 | 62.13 | 34 715 *(2.02%)* | 1 877 | 32 586 | 251 | 1 |
| 206SUR | 10 122 199 | 19.22 | 23 435 *(1.37%)* | 11 020 | 12 402 | 13 | 0 |
| 208SUR | 10 803 020 | 24.33 | 25 493 *(1.49%)* | 2 650 | 22 793 | 50 | 0 |
| 209SUR | 11 023 554 | 37.77 | 21 038 *(1.23%)* | 2 915 | 18 088 | 35 | 0 |
| 210SUR | 10 253 900 | 14.15 | 19 307 *(1.13%)* | 4 099 | 15 202 | 6 | 0 |

**Supplementary Table 2:** Number of variants for the 27 samples, total and separation according to the number of alleles found at the loci in each sample.

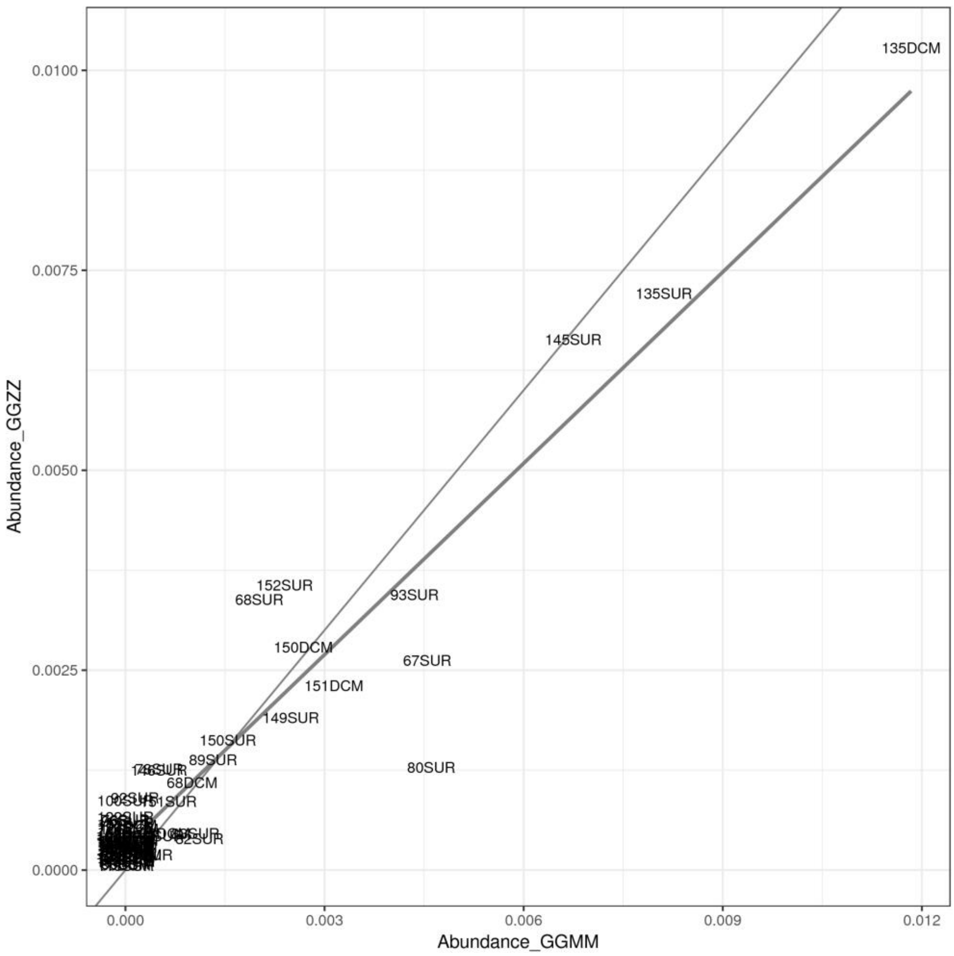


**Supplementary Figure 1:** Comparison of *Bathycoccus prasinos* RCC105 relative abundance in samples from the 0.8-5µm (X axis) and 0.8-2000µm (Y axis) size fractions in the same stations. The two lines correspond to the identity line (1:1 thick line) and to a linear regression from relative abundances values (thin line).

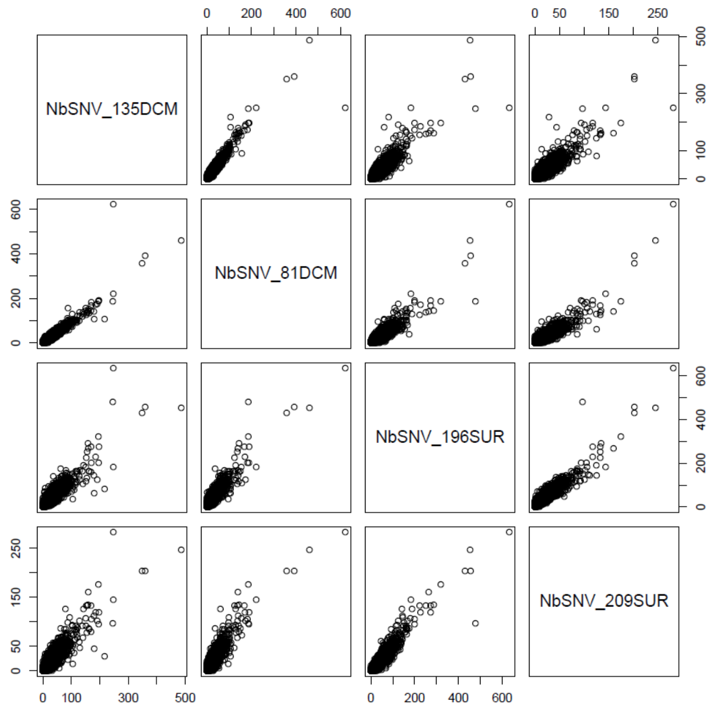


**Supplementary Figure 2:** Pairwise SNV number per gene in the four best-covered samples

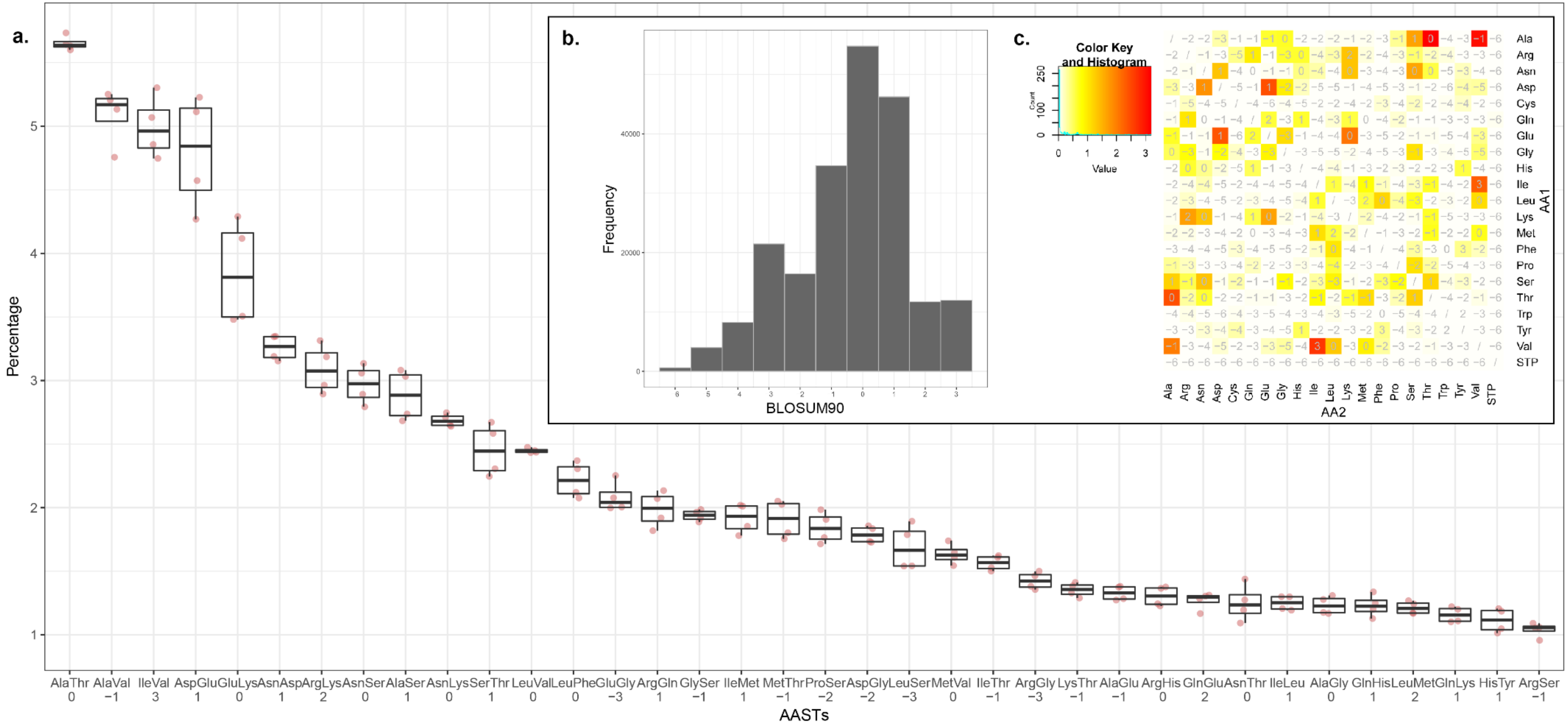

**Supplementary Figure 3:**

**a.** Frequencies of AASTs with more than 1% prevalence for the four best-covered samples. Associated BLOSUM90 scores are indicated under the amino-acid pair on the x-axis. Among 210 070 SAAVs, we observed 206 out of 210 theoretically possible AASTs and very similar frequencies among samples. The ten most common AASTs in all samples have an average score of 0.7, while the 10 rarest AASTs have an average score of -4.5

**b.** BLOSUM90 score distribution including all SAAVs, considering the transition between the two major amino-acids. The global distribution of all SAAV associated scores also shows a majority of positive values.

**c.** Percentage of pairs of amino-acids (AASTs) found in SAAVs. Colour corresponds to the average percentage between the 4 best-covered samples, ranging from white (absence) to red (high percentage). As expected, the most common and prevalent amino-acid substitutions would have low impact at the protein level given their positive BLOSUM90 substitution matrix scores.

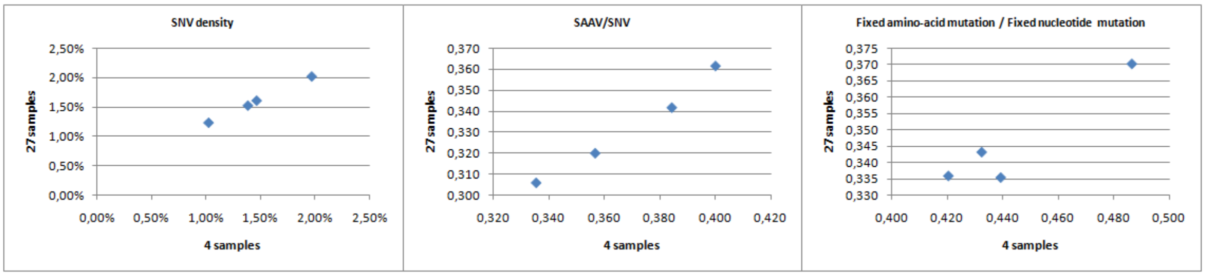


**Supplementary Figure 4:** Comparison of the SNV density, SAAV/SNV ratio, fixed amino-acid variant/fixed nucleotide variant ratio in the four samples when analysing them in the 4-sample set and the 27-sample set.

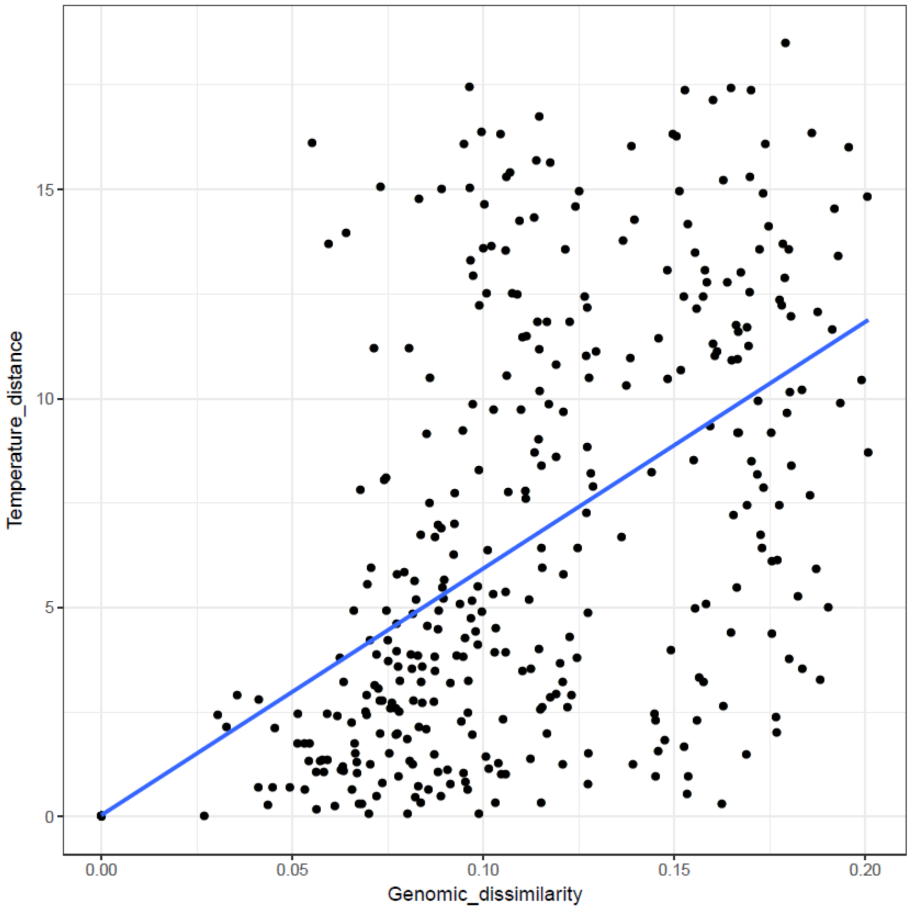


**Supplementary Figure 5:** Genomic dissimilarity (1 - genomic distance computed according to Methods) versus delta temperature (difference between two samples) for each pair of samples. The linear regression line is indicated in blue. Mantel test between the two parameters indicates a 0.4818 statistic at p-value=0.001.

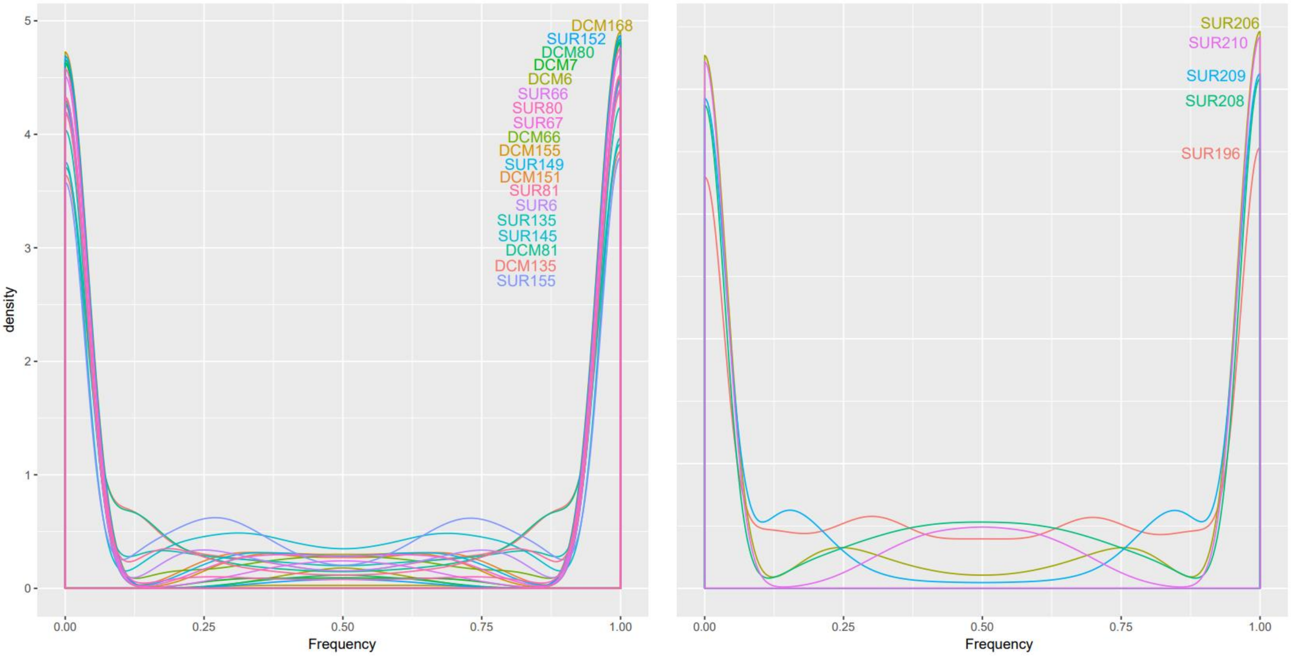


**Supplementary Figure 6:** Allele frequency distributions for all SNVs for different samples belonging to the temperate (left panel) or the arctic (right panel) cluster.

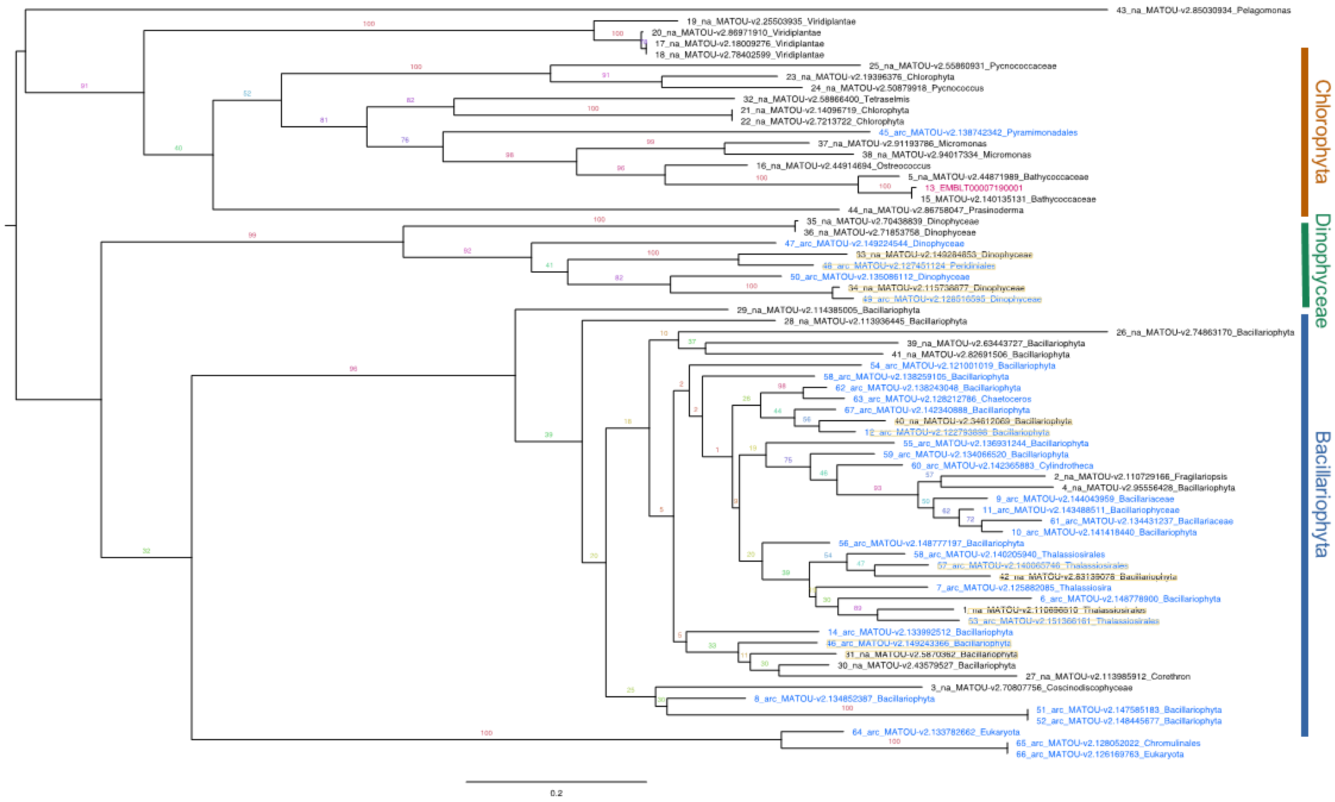


**Supplementary Figure 7:** Tree constructed on the basis of the alignment of mesophilic (black) and psychrophilic (blue) sequences of close eukaryotic EPSPS homologs found and sorted in Ocean Gene Atlas.

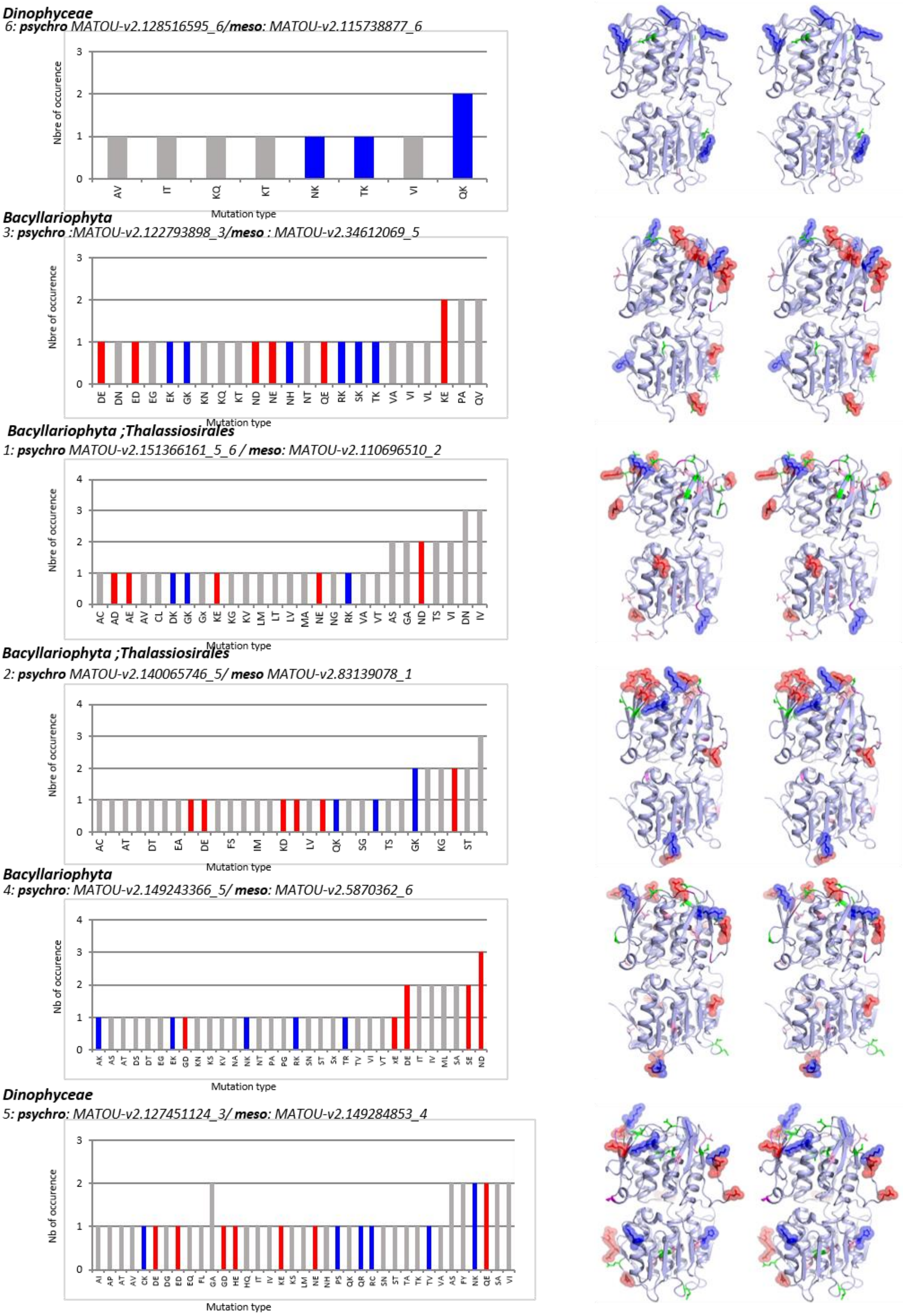


**Supplementary Figure 8:** (Left) Types and occurrences of mutations detected in the psychrophilic sequences compared to the corresponding mesophilic sequences for the 6 mesophilic/psychrophilic pairs selected in the tree in Supplementary Figure 8. They are arranged in ascending order of mutation number (top to bottom). (Right) Stereo views of the corresponding models showing the positions of charged residue mutations (positive: blue; negative: red).

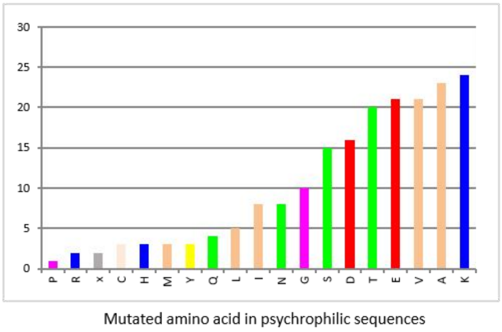


a

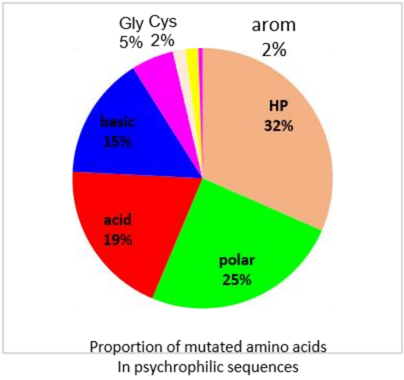


b

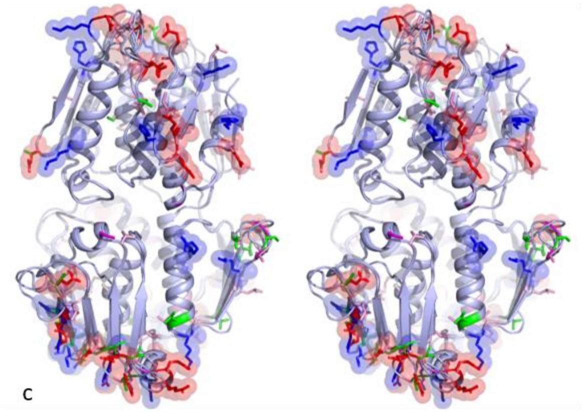

c

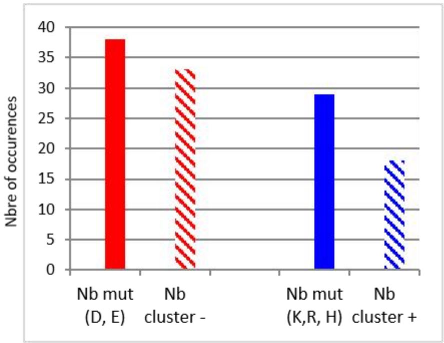


d

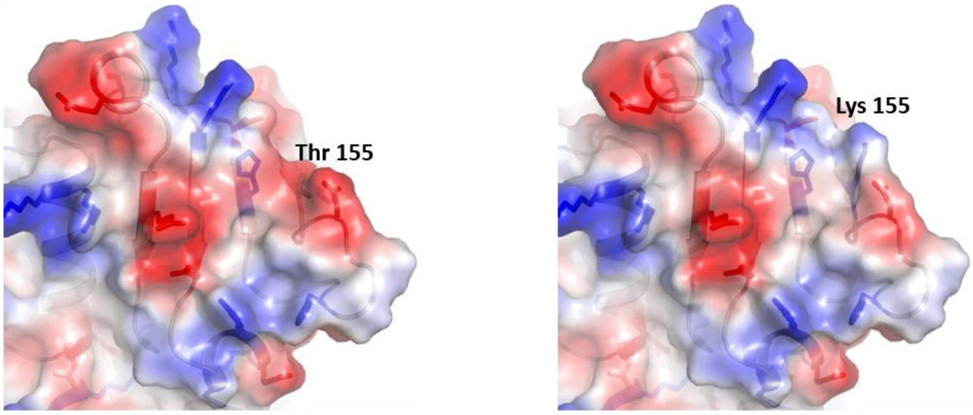


***meso**: MATOU-v2.115738877_6*

***psychro** MATOU-v2.128516595_6*

e

**Supplementary Figure 9:**

a: total number of mutated amino acids in the 6 selected psychrophilic EPSPS sequences.

b: Proportion of mutated amino acids per category for the total mutations observed in the six selected sequences in the Supplementary Figure 8 tree.

c: Structural superposition of the 6 psychrophilic EPSPS models (from Supplementary Figure 9) showing the location of all charged residue mutations with transparent spheres (red: negatively and blue: positively charged charged amino acids).

d: Total occurrence of charged residue mutations (solid rectangles) and charged residue mutations near similarly charged residues (striped rectangles) in the psychrophilic EPSPS structure models of the six sequences analysed. Red= glu, asp; blue=lys, arg or his.

e: comparison of electrostatic potentials of mesophilic (left) and psychrophilic (right) dinophyceae EPSPS models (first model in Fig. S9)
